# Applying Corrections in Single-Molecule FRET

**DOI:** 10.1101/083287

**Authors:** Antonino Ingargiola

**Affiliations:** Dept. Chem. & Biochem, Univ. California Los Angeles, Los Angeles, CA, USA

## Abstract

Single-molecule Förster resonance energy transfer (smFRET) experiments can detect the distance between a donor and an acceptor fluorophore on the 3-10nm scale. In ratiometric smFRET experiments, the FRET efficiency is estimated from the ratio of acceptor and total signal (donor + acceptor). An excitation scheme involving two alternating lasers (ALEX) is often employed to discriminate between singly– and doubly-labeled populations thanks to a second ratiometric parameter, the stoichiometry *S*. Accurate FRET and *S* estimations requires applying three well-known correction factors: donor emission leakage into the acceptor channel, acceptor direct excitation by the donor excitation laser and the “gamma factor” (i.e. correction for the imbalance between donor and acceptor signals due to different fluorophore’s quantum yields and photon detection effciencies).

Expressions to directly correct both raw FRET and S values have been reported in [1] in the context of freely-diffusing smFRET. Here we extend Lee *et al.* work providing several expressions for the direct excitation coeffcient and highlighting a clear interpretation in terms of physical parameters and experimental quantities. Moreover, we derive a more complete set of analytic expressions for correcting FRET and *S*. We aim to provide a clear and concise reference for different definitions of correction coeffcients and correction formulas valid for any smFRET experiment both in immobilized and freely-diffusing form.

## 1 Introduction

Förster resonance energy transfer (FRET) is a Coulombic interaction between the dipoles of two fluorophores, which results in the resonant and non-radiative transfer of excitation energy from a donor to an acceptor fluorophore (and the energy transfer probability decreases with the sixth power of the distance). Donor de-excitation via FRET competes with the donor’s intrinsic radiative and non-radiative de-excitation paths. Therefore, in the presence of a nearby acceptor, the lifetime of the donor is reduced. The quantum yield, or efficiency, of the FRET process can be computed as [2]:

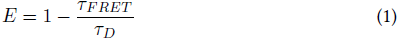

where *τ_FRET_* is the D lifetime in presence of FRET and *τ_D_* is the intrinsic D lifetime without any acceptor nearby. Computing *E* following eq. 1 requires measuring the D excited-state lifetime, for example using a TCSPC setup. A simpler method of estimating *E* consists in measuring only the intensity of donor and acceptor fluorescence (*F_D_* and *F_A_* respectively) and computing the FRET efficiency ratiometrically as:

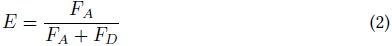

The previous eq. 1 and 2 require that the fluorescence lifetimes or intensities be relative to a single specie. In biological samples, where almost inevitably multiple FRET populations are present, single-molecule FRET (smFRET) experiments allows identifying different sub-populations and, for each of them, estimating the FRET efficiency [3].

The ratiometric approach of computing *E* is very common in smFRET owing to its modest hardware requirements (compared to TCSPC measurements) and has been extensively applied both to freely-diffusing and to surface-immobilized experiments. Unfortunately, unlike lifetime-based experiments, ratiometric FRET is affected by three systematic errors (or biases) intrinsic to the way *F_A_* and *F_D_* are measured. The first, a fraction of the donor emission spectrum almost inevitably falls in the acceptor detection band, causing spurious increase in acceptor-channel signal named “donor leakage”. Additionally, the acceptor signal is contaminated by a fraction of fluorescence due to direct excitation of the acceptor fluorophore by the donor laser (ideally the acceptor should only be excited by the donor). Finally, the relative (detected) donor and acceptor fluorescence intensity is biased because of the different fluorescence quantum yields and photon detection effciencies in the two detection channels (requiring the so-called “gamma factor” correction). These biases are well-known and expressions for their correction have been derived [1].

Contrary to ensemble measurements, single-molecule experiments can resolve different subpopulations and recover mean/peak FRET effciencies of each single conformational or binding state (at least in cases where there are no conformations that interconvert much faster than diffusion times). However, obtaining accurate mean FRET effciencies also requires applying corrections for the aforementioned biases.

This paper extends the ratiometric FRET corrections reported in Lee *et al*. [1]. We define the acceptor direct excitation as a function of different observable. For each definition we derive the direct excitation coeffcient as a function of physical parameters and discuss its physical interpretation. We also derive a complete set of formulas for computing *E* or *S* as a function of the raw *E* and *S* as well as of the aforementioned correction factors. Note that the expression here presented are valid for any ratiometric smFRET or ALEX-smFRET experiment, being it immobilized or freely diffusing [2, 3].

## 2 Definitions

### 2.1 Fluorescence intensities

We start by defining the fluorescence intensity signal as a function of the physical parameters. For surface-immobilized measurements the signal can be donor or acceptor counts acquired in a camera frame for a given fluorophore. For freely-diffusing experiments the signal can be the counts detected in the donor and acceptor channel during a “burst” (i.e. a single molecule crossing the excitation volume). Following [1] we define:

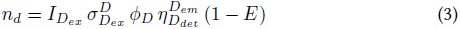

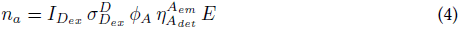

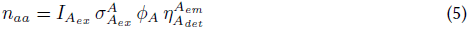

Eq. 3 and 5 are the detected quantities (e.g. counts, or camera intensity) after background correction in the DexDem and AexAem photon streams respectively. The *n_a_* quantity (eq. 4) needs to be estimated correcting the measured counts 
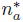
 in the DexAem stream (see eq. 10). The factors *I*, *σ*, *φ* and *η* are, respectively, the excitation intensity, the absorption cross-section, the fluorophore quantum yield and the photon detection efficiency. The label *D_ex_* (resp. *A_ex_*) indicates a coeffcient computed at the donor-laser excitation wavelength. *D_det_* (resp. *A_det_*) indicated the donor detection band. Finally, in the *σ* coeffcient the superscript D or A indicates the fluorophore. In addition to these quantities, we need to introduce the correction coeffcient *γ* and *β* which are defined as follows:

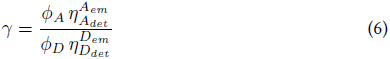

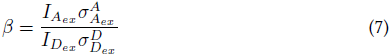

Briefly, *γ* makes the DexDem and DexAem signals commensurable (i.e. on the same scale) taking into account difference in dyes quantum yields and photon detectioneffciencies. Similarly, the *β* factor is used to make the total Dex signal commensurable with the AexAem signal by taking into account the differences in excitation intensities (*I_A_ex__* vs *I_D_ex__*) and in dyes absorption cross-sections 
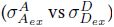
. This expression of the *β* coeffcient has been derived in [1] during the derivation of the fitting procedure for *γ*-factor.

It is also useful to introduce the total corrected signal during D-excitation which we can define equivalently with one of the following expressions:

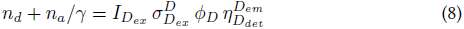

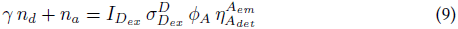

The choice between eq. 8 and 9 is only matter of convention. Finally, in a real experiment we cannot measure *n_a_* directly, instead we acquire a value 
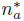
 that is contaminated by donor leakage (*Lk*) and acceptor direct excitation (*Dir*). We define 
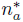
 and the correction terms as follows:

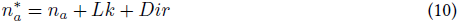

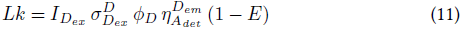

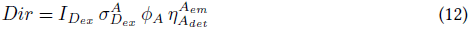

Consistently with the *n_d_* and *n_aa_* definitions (eq. 3 and 5), the quantity 
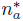
 is assumed already background corrected.

### 2.2 FRET and Stoichiometry

We start defining the FRET effciency *E* and the proximity ratios *E_PR_* and *E_R_*:

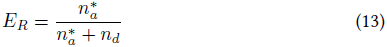

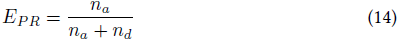

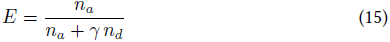

where *n_d_*, *n_a_* are the donor and acceptor detected counts after all the corrections (see eq. 3 and 4), while 
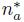
 are the acceptor counts with only background correction of eq. 10 (no leakage and direct excitation corrections).

Similarly, for the stoichiometric ratio we can have different definitions depending on the degree on corrections that are applied:

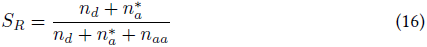

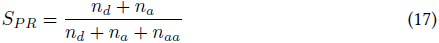

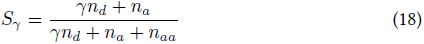

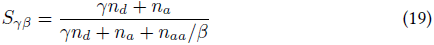

*S_R_* (eq. 16) is the raw stoichiometry without any correction except for background (see definition of 
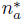
 in eq. 10). *S_PR_* (eq. 17) is the stoichiometry corrected for leakage and direct excitation (see *n_a_* definition in eq. 4). *S_γ_* (eq. 18) is the stoichiometric ratio corrected for leakage, direct excitation and *γ* (so that FRET populations have stoichiometry centered around a constant value, typically close to 0.5). *S_γβ_* (eq. 19) includes a *β* correction ensuring that FRET populations have stoichiometry centered around 0.5. Since *β* (eq. 7) is equal to the ratio 
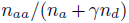
 (eq. 4 and 9) it follows that:

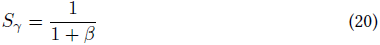

From eq. 20 follows that when *β* is known it is possible to compute the *S_γ_* value around which all FRET populations are distributed. As noted before, for *S_γβ_* this value is always 0.5.

### 2.3 Definition of direct excitation

The term *Dir* can be equivalently expressed as a fraction of any fluorescence intensity components (i.e. counts in the donor or acceptor channel during donor or acceptor excitation). Here we present five different definitions and their physical interpretation.

#### Definition 1

Defining *Dir* as a function of *n_aa_* we have:

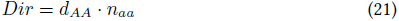

The coeffcient *d_AA_* can be computed from an acceptor-only population in ALEX measurement, because in this case eq. 10 becomes 
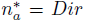
. In terms of physical parameters, recalling eq. 12 and 5, we can express *d_AA_* as:

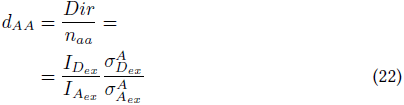

Since computing *Dir* through *d_AA_* requires the spectroscopic quantity *n_aa_* (see eq. 5), it cannot be used in case of single-excitation measurements. In this case the definitions in the next section can be used. Note that *d_AA_* is indicated as *d* in [1].

#### Definition 2

Defining *Dir* as a function of the “corrected total signal” as defined in eq. 8 ((*n_a_* + *γn_d_*)) results in:

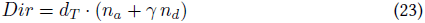

From eq. 23, it follows that:

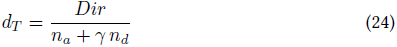

To derive the expression of *d_T_* as a function of physical parameters, consider the case of 100% FRET molecule. In this case, knowing that *n_d_* = 0 and recalling the expression of *n_a_* from eq. 4, we obtain:

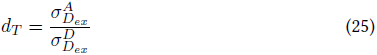

Noting that, for *E* < 1 the “corrected total signal” *n_a_* + *γn_d_* (e.g. the corrected burst size in freely-diffusing measurements) will not change when *γ* is constant. Therefore the previous expression is valid for any *E*.

In ALEX measurements is easier to estimate *d_AA_* from the data. Therefore, expressing *d_T_* as a function of *d_AA_* allows to easily estimate the former coeffcient from the data. From the definitions of eq. 22, 25 and 7 we obtain:

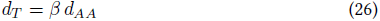

This relation follows from the definition of *β* reported in the previous section and originally defined in [1].

#### Definition 3

Defining *Dir* as a function of the “corrected total signal” as defined in eq. 9 (*n_a_/γ*+ *n_d_*) we have:

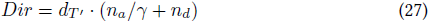

The coeffcient *d_T′_* can be obtained from the *d_T_* expression noting that we simply divide the “corrected total signal” by *γ*:

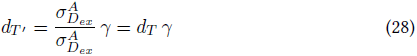

The coeffcient *d_T′_* is indicated as *d′* in [1] (main text p. 2943 and SI). Note that the definition of *d′* given in eq. (27) of [1] has been derived for a *E* = 0 population (for which *n_d_* + *n_a_/γ* = *n_d_*). However, by using the corrected total signal, it is possible to use the same coeffcient to express the *Dir* contribution for any FRET population (and independently from *E*).

#### Definition 4

Defining *Dir* as a function of *n_d_* we have:

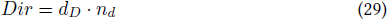

The coeffcient *d_D_* is a function of *E* as well as the physical parameters. Taking the ratio of the physical definitions of *Dir* and *n_d_* we obtain:

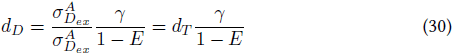

#### Definition 5

Defining *Dir* as a function of *n_a_*:

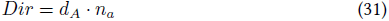

The coeffcient *d_A_* is a function of *E* as well as the physical parameters. Taking the ratio of the physical definitions of *Dir* and *n_a_* we obtain:

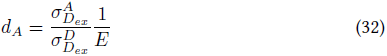

### 2.4 Discussion of Definitions 1-5

Definitions 4 and 5 are inconvenient because the coeffcient depends on *E*. Definition 3 does not depend on *E* but depends on *γ*, while Definition 2 depends only on the ratio of two absorption cross sections and is therefore the most general form. Definition 1 can only be used in an ALEX measurement but it is easy to fit from the *S* value of the A-only population.

So, for non-ALEX measurements, Definition 2 (*d_T_*) gives the simplest and most general coeffcient. It can be computed from datasheet values or from *d_AA_* estimated from an ALEX measurement using the same dyes pair and D-excitation wavelength (*d_T_* = *βd_AA_*).

As physical interpretation, definitions 2 and 3 are similar. In Definition 2, when *E* = 1, the “corrected total signal” is *n_a_*. When *E* < 1, the “corrected total signal” does not change (at the same excitation intensity, and fixed *γ*) being the sum of acceptor and *γ*-corrected donor counts. Similar considerations hold for Definition 3 (starting from *E* = 0). Note that using eq. 25 to estimate *Dir* requires the knowledge of the corrected total signal of eq. 8 (including A-direct excitation correction). For practical purposes, using a signal only corrected for *γ* and leakage to compute *Dir* via eq. 25, is a very good approximation. Alternatively, using eq. 33 (see next section) it is possible to compute corrected *E* values without any approximation.

### 2.5 Correction formulas

We can expressing *E* as a function of *E_R_* and the three correction factors as follows:

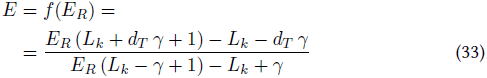

This expression is the same of eq. S9 in [1] when we replace *d_T_ γ* with *d′*.

Similarly we can express *S* as a function of *S_R_*, but in this case the expression will also depend on *E_R_* in addition of the correction parameters:

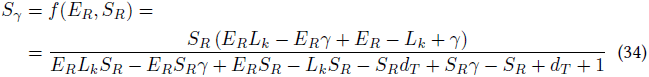

A similar formula has been reported in [1] (SI) expressing *S* as a function of *S_PR_* and *E_PR_*. Here the expression is simply expanded as a function of *E_R_* and *S_R_*, resulting in an explicit dependence on *lk* and *d_T_*. The derivation of these formulas only involves using algebraic manipulations of the *E* and *S* expression. To avoid trivial errors, these expression have been derived with computer-assisted algebra (CAS). We also provide text-based version of the formula (python syntax) that is tested and easy to copy and paste in most other text-based language. For derivation details see Appendix: Derivation of the formulas.

## 3 Conclusion

We have introduced five definitions of acceptor direct excitation as a function of different experimental observable, and discussed that out of the five, two have the most useful in practice. In particular, eq. 25 can be used to correct for A-direct excitation even in single-laser measurements provided the coeffcient *d_T_* can be estimated independently. Furthermore, eq. 33 and 34 allows to apply corrections to *E* and *S* values, only knowing the raw *E* and *S* and the correction factors. With eq. 33 and 34 it is possible to correct the fitted *E* or *S* values as a last independent step of the analysis, without the need to modify (i.e. correct) the distributions prior fitting. This is important because, from a statistical point of view, the fit of the raw *E* and *S* peaks can provide more reliable estimates due to simpler modeling (e.g. using a Binomial distribution) which requires less assumptions. For example, methods such as shot-noise [4] and probability distribution analysis [5,6] and Gopich-Szabo likelihood analysis [7,8] can be directly applied to raw FRET distributions. Conversely, applying these methods to the corrected FRET distributions requires unnecessary complex statistical models which include the effect of each correction factor. In practice, the benefit of a more complex model is dwarfed by the inaccuracies arising from the additional approximations (even implicit) and from reliance on estimated correction parameters in the model itself. Using eq. 33 and 34, instead, allow to decouple the correction of *E* and *S* values from the population-level statistical modeling, resulting in more robust models and more accurate estimates.

## Acknowledgments

We thank Dr. Eitan Lerner for a critical reading of the manuscript. Support provided by NIH R01 GM095904.

## Appendix: Derivation of the formulas

See Derivation of FRET and S correction formulas.

